# A High-throughput Neurohistological Pipeline for Brain-Wide Mesoscale Connectivity Mapping of the Common Marmoset

**DOI:** 10.1101/315804

**Authors:** Meng Kuan Lin, Yeonsook Shin Takahashi, Bing-Xing Huo, Mitsutoshi Hanada, Jaimi Nagashima, Junichi Hata, Alexander S. Tolpygo, Keerthi Ram, Brian Lee, Michael Miller, Marcello G.P. Rosa, Erika Sasaki, Atsushi Iriki, Hideyuki Okano, Partha P. Mitra

## Abstract

Understanding the connectivity architecture of entire vertebrate brains is a fundamental but difficult task. MRI based methods offer whole brain coverage, but remain indirect in the approach to connectivity mapping. Recent progress has been made in directly mapping whole-brain connectivity architecture in the mouse at the mesoscopic scale. The basic approach uses tracer injections systematically placed on a grid of locations spanning the brain and computational analysis of the resulting whole brain data sets. Scaling this approach to bigger primate brains poses nontrivial technical challenges. Here we present an integrated neurohistological pipeline as well as a grid-based tracer injection strategy for systematic mesoscale connectivity mapping in the common Marmoset (*Callithrix jacchus*). Individual brains are sectioned into ∼1700 20µm sections using the tape transfer technique, permitting high quality 3D reconstruction of a series of histochemical stains (Nissl, myelin) interleaved with tracer labelled sections. Combining the resulting 3D volumes, containing informative cytoarchitectonic markers, with *in-vivo* and *ex-vivo* MRI, and using an integrated computational pipeline, we are able to overcome the significant individual variation exhibited by Marmosets to obtain routine and high quality maps to a common atlas framework. This will facilitate the systematic assembly of a mesoscale connectivity matrix together with unprecedented 3D reconstructions of brain-wide projection patterns in a primate brain. While component instruments or protocols may be available from previous work, we believe that this is the first detailed systems-level presentation of the methodology required for high-throughput neuroanatomy in a model primate.

## 1 Introduction

The connectional architecture of the brain underlies all the nervous system functions, yet our knowledge of detailed brain neural connectivity falls largely behind genomics and behavioral studies in humans and in model research species such as rodents ^1^. To fill this critical gap, a coherent approach for the mapping of whole-brain neural circuits at the mesoscale using standardized methodology was proposed in 2009 ^1^. Since then, several systematic brain connectivity mapping projects for the mouse have been initialized and established, including the Mouse Brain Architecture Project ^2^ (www.brainarchitecture.org), the Allen Mouse Brain Connectivity Atlas ^3^ (connectivity.brain-map.org), and the Mouse Connectome Project (www.mouseconnectome.org). Non-human primates (NHPs) were also proposed as an important group in which to study whole-brain neural architecture. However, the high-throughput experimental approaches for mouse do not automatically apply to NHPs due to bioethical as well as experimental considerations, larger brain sizes coupled with stringent limitations on the numbers, as well as limitations arising from the increased individual variability of the brains.

There has been an increase in the usage of the common marmoset (*Callithrix jacchus*) as a model organism in contemporary neuroscience research^4-8^ (Supplementary Figure 1). Marmosets offer a number of experimental advantages, including lower cost, ease of handling and breeding ^5,9^, smaller brain sizes (≈35mm*25mm*20mm) that potentially allow more comprehensive analysis of the neuronal circuitry, and importantly the development of transgenic marmosets and the application of modern molecular tools ^10-12^.

Marmosets are New World monkeys, in contrast with the Old World Macaque monkeys which are the pre-eminent NHP models used in basic and pre-clinical neuroscience research. As depicted in Figure 1a, New World monkeys, together with Old World monkeys, apes and humans, form the simian primates (order Primates, infraorder Simiiformes). Simians diverged from prosimians such as lemurs and lorises approximately 85 million years ago (Mya). Among the simians, New World monkeys have evolved in isolation from Old World monkeys, apes and humans for at least 40 million years. Prima facie this seems to indicate a relative weakness in using Marmosets as NHP models in contrast with the Macaques. Nevertheless, a good case can be made for Marmosets as NHP models of humans, despite the earlier evolutionary divergence.

Marmosets exhibit more developed social behavior ^13^ and vocal communication ^14^ traits, thus social-vocal Human traits (and corresponding dysfunctions) are better modeled in Marmosets than in Macaques. This is either the product of convergent evolution, or are shared ancestral traits preserved in the Human lineage (but lost in the Macaque due to selective niche adaptation). Marmoset brains are smaller than Macaque brains and are comparable in size to some rodents (cf. Squirrels and Capybara, both species of rodents, have brain volumes comparable to Marmosets and Macaques). However Marmosets are phylogenetically closer to Humans than Rodents, and thus have more commonality in terms of brain architecture (proportionately larger and more differentiated higher order cortical areas, as opposed to primary cortical areas ^15^ (Figure.1).

**Figure 1.**
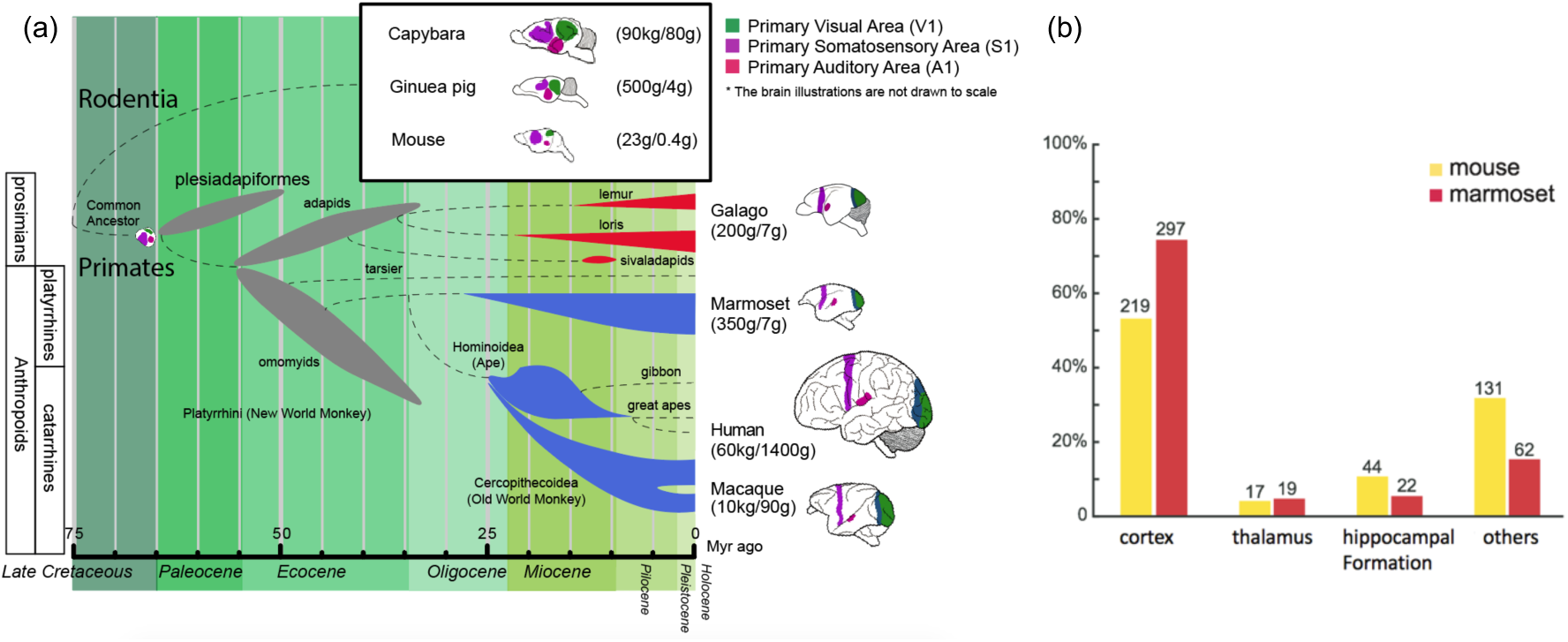
(a) Phylogenetic tree ^16-23^ showing the ancestral history of extinct and extant primates, after divergence from the common ancestor with rodents (top right inset box) *ca.* 75 million years (Myr) ago. The bottom bar shows geological eras. Thickness of spindle shaped areas in the evolutionary tree indicate of prosperity (estimated population and numbers of species) of the group along the history in extinct (gray) prosimian (red) and simian (blue) primates. Each bifurcation represents the species divergence, although the divergence time typically has a wide range and remains controversial. Primates diverged into platyrrhini, the New World Monkey, and catarrini, around 38.9-56.5 million years ago. Catarrini further evolved into Ape, including humans, and Old World Monkey as well as macaque monkeys 25.1-37.7 million years ago. Sketches of the brain in each species are shown on the right, next to their species name. The colored areas in the various brain illustrations indicate the primary visual area as green, somatosensory as purple, and auditory areas as red; each represents an extant primate (bottom right row) and rodent (top inset box) species’ body weight (first numbers in brackets) and brain weight (last numbers in brackets) sizes ^24-26^. Phylogenetic tree adapted from Masanaru Takai ^27^. (b) Fractional brain region volumes, and numbers of injection sites used in grid-based injection plans for marmoset ^28^ and mouse ^29^. Bar plots show the relative compartment volumes as percentages of total grey matter volume. The numbers on top of the bars show the number of injection sites within the region in each species. The grids correspond to spacing between injection sites of ∼1 mm isometric in mice, and ∼2-3 mm isometric in marmosets.

Following the BRAIN (Brain Research through Advancing Innovative Neurotechnologies) Initiative in the U.S. and the HBP (Human Brain Project) in Europe in 2013, Japan launched the Brain/MINDS project (Brain Mapping by Integrated Neurotechnologies of Disease Studies) with a focus on the common marmoset (*Callithrix jacchus*) as an NHP model ^4^ (http://www.brainminds.jp/). As part of Brain/MINDS, a combined histological/computational pipeline was established at RIKEN to develop a mesoscopic whole-brain connectivity map in the Marmoset. The corresponding methodology is described in this manuscript.

Tract-tracing methods remain the gold standard for studying neural circuit structure at the whole brain level ^30^. Previous brain-wide connectivity mapping for non-human primates have been based on literature curation and meta-analyses. A pioneering survey by Felleman & Van Essen ^31^ reviewed 52 studies, including both anterograde and retrograde tracing results, to generate a connectivity matrix of 33 brain regions in the visual system of macaque monkeys (Table 1). Building upon Felleman & Van Essen ^31^, a more comprehensive database of macaque brain connectivity, CoCoMac (Collation of Connectivity data on the Macaque brain, cocomac.g-node.org) ^30,32,33^, surveyed over 400 tracing studies with ∼3,300 injections and established a connectivity matrix of 58 brain regions ^34,35^ (Table 1). While the historical tracing studies mostly contain qualitative information, more recent studies have aimed at building a quantitative connectivity database of the macaque brain ^36-38^ (core-nets.org; Table 1).

**Table 1.**
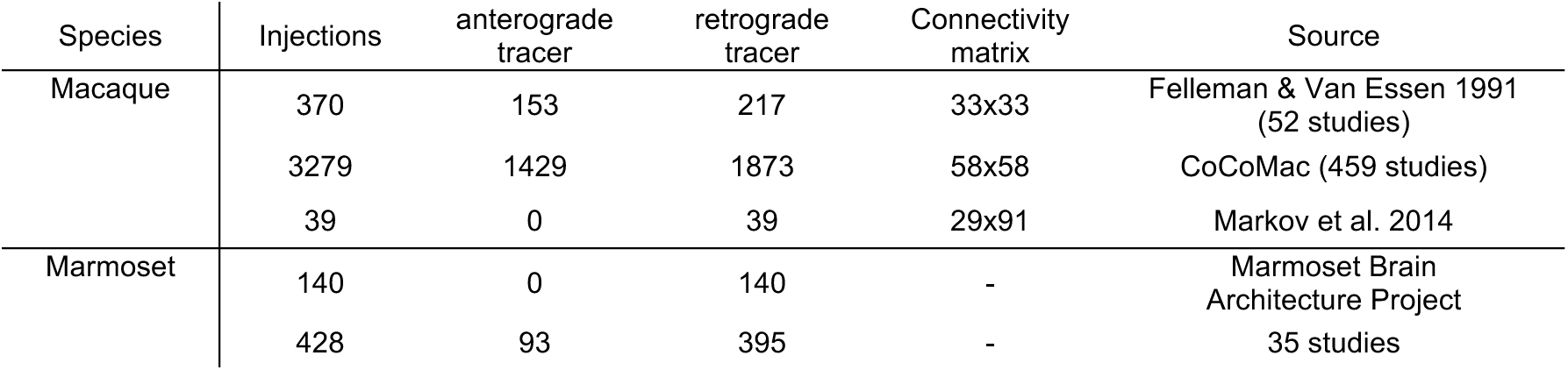
Summary of historical tract-tracing studies in macaque and marmoset monkeys. Three resources of macaque monkeys brain connectivity are shown. Felleman & Van Essen ^31^ and CoCoMac each surveyed a set of studies to generate the connectivity matrix (full reference list in Supplementary reference list). Note that CoCoMac is inclusive of the work collected in Felleman & Van Essen ^31^. Around 235 injections lackf tracer direction information. Markov et al. 2014 ^36^ was a single study using only the retrograde tracer to generate the connectivity matrix as well as quantifying the connection strengths. To date the most comprehensive marmoset brain connectivity resource available online includes 140 retrograde tracing studies (http://marmoset.braincircuits.org/injection). A complete connectivity matrix is not yet available for the marmoset brain. We have ourselves surveyed 35 marmoset brain tracing studies that contain 428 tracer injections including both anterograde and retrograde tracers. For both macaque and marmoset brain injections, bidirectional tracer injections were double counted as one anterograde and one retrograde tracer injection.

For the Marmoset, an online database of >140 retrograde tracer injection studies in about 40 brain regions is available online (http://marmoset.brainarchitecture.org) ^39^. By surveying 35 tract tracing studies (Supplementary reference list) in marmosets since the 1970s, we have collected data from over 400 injections, but much of this knowledge cannot be easily integrated with current efforts given the use of older nomenclatures, and the lack of access to primary data. A full connectivity matrix is yet to be established (Table 1). Nevertheless existing knowledge about the marmoset visual, auditory, and motor systems indicate strong similarities between marmoset and macaque brain circuitry, suggesting a preserved brain connectivity plan across primates ^40-42^. Comparing two NHP brain architectures (Marmoset, Macaque) will help to better contextualize Human brain circuit architecture.

None of these earlier studies in NHPs have used a single, consistent methodology employing a unified experimental-computational workflow, dedicated to systematic mesoscale connectivity mapping. This is the goal of the pipeline described in this paper. Importantly, brain-wide data sets are already available for grid-based tracer mapping projects in the Mouse. A corresponding data set generated using similar techniques will allow us to gain a more unified view of primate brain connectivity architecture, and also permit an unprecedented comparative analysis of mesoscale connectivity in Rodents and Primates.

## 2 Methods

A high throughput neurohistological pipeline was established at the RIKEN Center for Brain Science, based on the pipeline developed for the MBA project ^2^ at CSHL. The pipeline employed a customized tape-transfer assisted cryo-sectioning technique to preserve the geometry of individual sections. Each brain was sectioned serially into a successive series of four 20um sections: a Nissl stained section, a Silver (Gallyas) myelin stained section, a section stained (ABC-DAB) for the injected cholera toxin subunit B (CTB) tracer and an unstained section imaged using epifluorescence microscopy to visualize the results of fluorescent tracer injections. Three types of fluorescent neural tracers were injected into the brain to reveal the mesoscale neural connectivity. The four sets of sections: Nissl, myelin, CTB and fluorescent sections were processed and imaged separately, and later re-assembled computationally. A computational pipeline was established to perform high-throughput image processing. A common reference atlas ^43,44^ was registered to each individually reconstructed brain series and the projection strengths were suitably quantified.

### 2.1 Experimental pipeline

All experimental procedures were approved by the Institutional Animal Care and Use Committee at RIKEN and a field work license from Monash University, and conducted in accordance with the Guidelines for Conducting Animal Experiments at RIKEN Center for Brain Science and the Australian Code of Practice for the Care and Use of Animals for Scientific Purposes. Female marmosets (*Callithrix jacchus*), 4 to 8 years old, 330g - 440g in weight, were acquired from the Japanese Central Institute for Experimental Animals.

#### *In-vivo* MRI

Upon habituation, the marmosets promptly went through magnetic resonance (MR) imaging. MR scans were performed using a 9.4T BioSpec 94/30 US/R MRI scanner (Bruker, Biospin, Ettlingen, Germany) with actively shielded gradients that had a maximum strength of 660 mT/m. Several MRI protocols were carried out for each individual marmoset. T1 mapping and T2-weighted images (T2WI) were used in *in-vivo* MR imaging. More details of the scan protocol can be found in Supplementary Information A.

#### Neuronal tracer injections

For parsimony, four tracers were placed in the right hemisphere of each marmoset, including two anterograde tracers: AAV-TRE3-tdTomato (AAV-tdTOM) and AAV-TRE3-Clover (AAV-GFP), and two retrograde tracers: Fast Blue (FB) and CTB. Surgical procedures for tracer injections were adapted from the previously established protocols^45-47^. Tracers were delivered at the injection sites using Nanoject II (Drummond, USA), with dosage controlled by Micro4 (WPI, USA). For cortical injections, each tracer was delivered with depths of 1200µm, 800µm, and 400µm sequentially perpendicular to the cortical sheet, with equal volumes. The planning for tracer injections approximately followed a uniform 2 ×; 2 ×; 2mm grid spacing, intended to cover the entire brain cortical and subcortical regions^48^ (Supplementary Information B). The current data set used to validate the method presented here includes 118 injections. At each injection site, one retrograde and one anterograde tracer was injected in separate to cover the efferent and afferent projections of that site. Figure 2a,b shows currently covered injection sites.

#### *Ex-vivo* MRI and cryo-sectioning

After tracer injection and a 4-week incubation period, the marmoset brain was perfused with a 0.1M phosphate buffer (PB) flush solution followed by 4% paraformaldehyde (PFA) in 0.1M PB fixation solution. The same MR scan protocol for *in-vivo* MRI was used for *ex-vivo* Diffusion Tensor Imaging (DTI) scanning. Additional high-resolution (300µm) T2-weighted images (T2WI) were carried out for *ex-vivo* MR imaging (Supplementary Information A). Following fixation, the brain was transferred to 0.1M PB to take an *ex-vivo* MRI. It was then immersed in 10% then 30% sucrose solution over a 48-hour period to safeguard against thermal damage. The brain was embedded in freezing agent (Neg-50^TM^, Thermo Scientific 6505 Richard-Allan Scientific) using a custom developed apparatus and a negative cast mold of the brain profile. The apparatus was submerged in an optimal cutting temperature compound to expedite the freezing process ^49^. More details can be found in Supplementary Information C.

Cryo-sectioning of the brain was performed using a Leica CM3050 S Cryostat in a humidity chamber set at 18°C and 80% humidity. The cryostat specimen temperature was set to −15 to −17°C while the chamber temperature was set to −24°C. This temperature differential was used to make certain the tissue was never in danger of being heated unnecessarily. Brains were cryo-sectioned coronally on a custom made cryostat stage using the tape transfer and UV exposure method^2^ (Supplementary Information D). Every four consecutive sections were separately transferred to four adjacent slides, to establish the four series of brain sections to be stained for different methods. Each section was 20µm in thickness, hence the spacing between every two consecutive sections in the same series was 80µm. The four slides were transferred and cured for 12 seconds in a UV-LED station within the cryostat. All cured slides were placed inside a 4°C refrigerator for 24 hours to allow thermal equilibration.

#### Histological staining

Separate histological staining processes were performed on the different series of brain sections (Supplementary Information E). High-throughput Nissl staining of neuron somata was performed in an automated staining machine (Sakura Tissue-Tek Prisma, DRS-Prisma-J0S) (Figure 2c). The myelin staining technique used a modified ammoniacal silver stain originally developed by Gallyas^50^. The present modification provided higher resolution of fiber details that could be used for myeloarchitecture identification. A representative magnified image of myelin staining in the V4 (middle temporal crescent) visual cortex is shown in Figure 2d. Using a modified protocol developed for the MBA project at CSHL, the staining of retrograde and anterograde CTB label was successfully attained ^51^ (Figure 2e). Finally, retrograde fluorescent tracers revealed originating somata while the anterograde tracers revealed projecting axons from fluorescent imaging. Figure 2(f-h) shows the simultaneous fluorescent tract tracing using AAV-GFP, AAV-tdTOM and FB within the same brain. More detailed high-magnification images can be found in Supplementary Figure 2.

**Figure 2.**
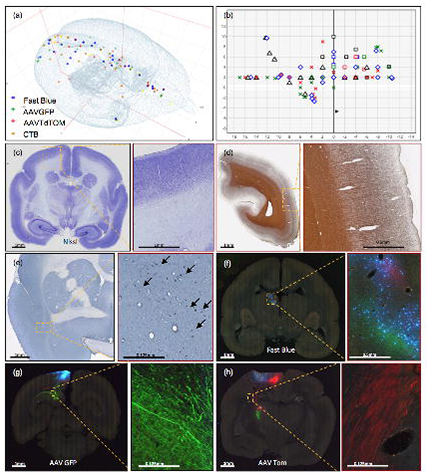
(a, b) Current injection sites in the marmoset cortex in (a) 3D and (b) 2D dorsal view the cortex only, in stereotaxic coordinates ^43^. The grid in (b) shows current successful injections. Each tracer is represented with a different color of marker: blue: Fast Blue; green: AAV-GFP; red: AAV-tdTOM; brown: CTB. Two tracers, one anterograde and one retrograde, are injected at each site. (c-h) Sample coronal brain section images of four series. (c) A coronal section after Nissl staining is shown with increasing magnification. Around Area 3a (magnification box), 6 cortical layers and the white matter are clearly differentiable based on cell body density. (d) A coronal section of the left hemisphere after silver staining showing myelin. Around Visual area V4T (Middle Temporal) crescent; magnification box), layers I-VI can be clearly characterized based on the myelin fiber density. Heavy myelination can be seen in layer 3 and continues into layer 4-6 with clear inner and outer bands of Baillarger. (e) Partial coronal section after immunohistochemistry treatment for CTB. After injection into Area 10, CTB labeled neurons were found in the claustrum (magnification box). The arrows indicate CTB-labeled cells at 0.125mm. (f-h) Coronal sections in different parts of the brain showing fluorescent tracers including (f) retrograde tracer Fast Blue (g) anterograde tracer AAV-GFP, and (h) anterograde tracer AAV-tdTOM.

The pipeline adopted the Sakura Tissue-Tek Prisma system for high-throughput staining purposes. Upon completion of auto staining, the system loaded the dehydrated slides into an automatic coverslipper (Sakura Tissue-Tek Glas, GLAS-g2-S0) where 24 ×; 60mm cover glass (Matsunami, CP24601) were applied with DPX mounting media (Sigma, 06522); then put into drying racks for 24 hours. Figure 3 shows the overall steps as well as time taken to process one marmoset brain before moving to the computational pipeline starting with imaging.

**Figure 3.**
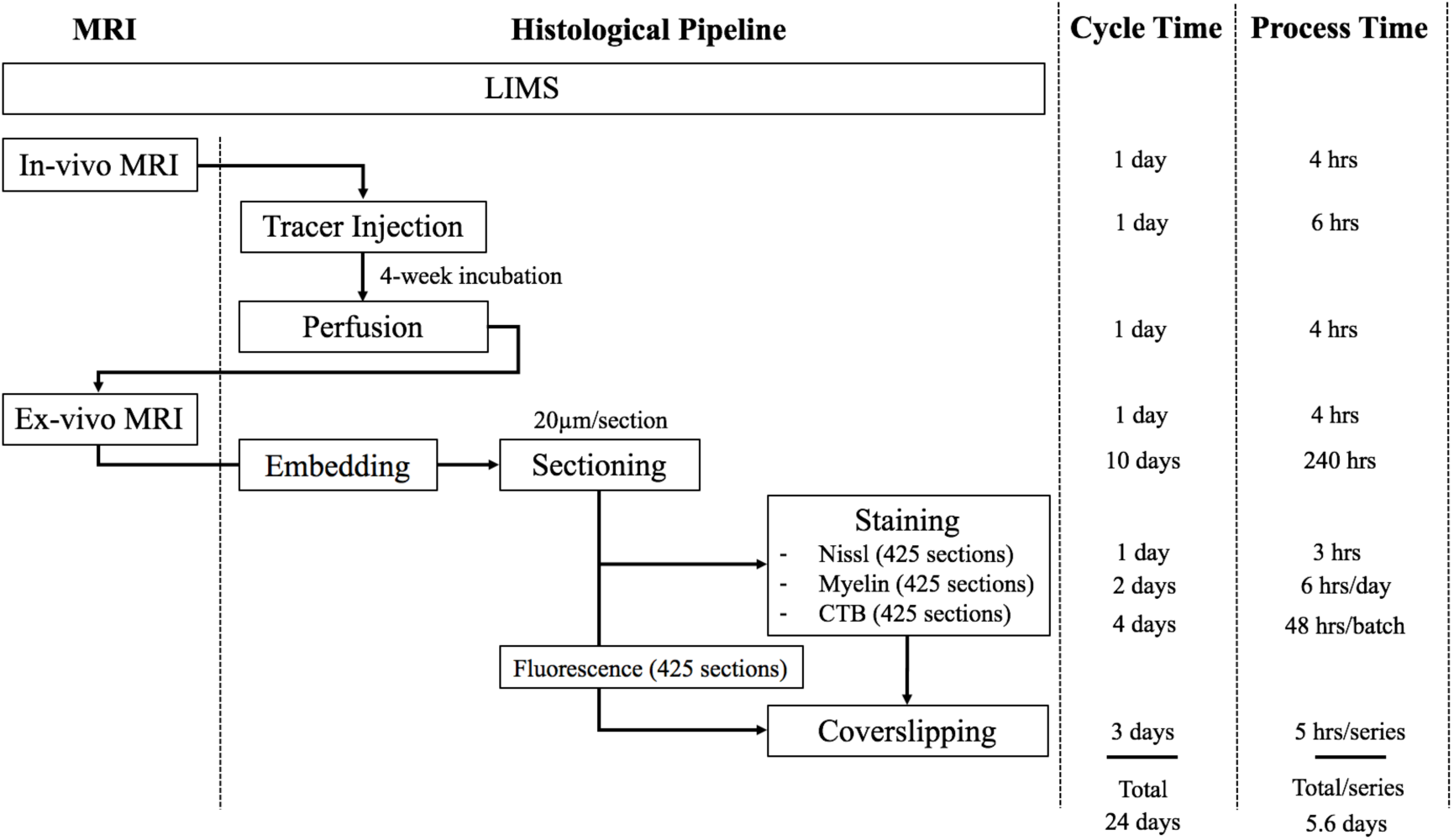
The workflow of the experimental pipeline and the processing time for one marmoset brain. Arrows show the sequence of individual experiments. A custom-made LIMS (Laboratory Information Management System) performs housekeeping for the entire process and constitutes an electronic laboratory notebook. The entire brain is sectioned into ∼1700 sections, ∼ 400 in each series. Each series include ∼295 slides, comprising of 1/3 of the slides with 2 brain sections mounted and 2/3 with 1 brain section/slide. Coverslipping includes the drying and clearing stages. The processing time does not include the overnight waiting period after sectioning in each batch. The overnight incubation time is excluded in the CTB procedure as well as the overnight dehydration in a myelin stain. Processing Time on the right shows the time involved in processing each experimental step, in hours. The Cycle Time (in days) shows the total time required to initiate and finish the entire procedure from start to finish, including quiescent periods, before commencing the procedure for another brain. Total time on the bottom is not a summation of the individual procedure times above because of parallel, pipelined processing which reduces total processing times. For example, when Nissl series are being processed in the automatic tissue staining machine for Nissls, CTB and myelin staining can be performed simultaneously at other workstations.

Including imaging, one full Nissl brain series can be completed in 6 days. The myelin series including imaging requires 6.4 days using a limited 60-slide staining rack. The CTB series took a total of 7.9 days to complete due to batch limitations (3.5 batches with 120 slides/batch in total). The time for completion for the fluorescent brain series was 8 days; the slide scanning time on the Nanozoomer used in the project is approximately twice the brightfield scanning time. Overall, the four separate series of one brain could complete in two weeks (a pipeline processing rate can be found in Supplementary Information H). The digitized brains are then passed onto the computational pipeline including atlas registration, cell and process detection and online presentation.

### 2.2 Computational pipeline

All the prepared slides were scanned by series with a Nanozoomer 2.0 HT (Hamamatsu, Japan) using a 20x objective (0.46 µm/pixel in plane) at 12-bit depth and saved in an uncompressed RAW format. Nissl, myelin and CTB series were brightfield scanned. Fluorescence series were scanned using a tri-pass filter cube (FITC/TX-RED/DAPI) to acquire the 3 RGB color channels for each slide. A Lumen Dynamics X-Cite *exacte* light source was used to produce the excitation fluorescence.

The RAW images for all four series of slides comprise ∼9 terabytes of data for each brain. In order to process these large data volumes, the pipeline includes networked workstations for data-acquisition, image processing and web presentations. All systems were connected to two directly attached data storage nodes to ensure that all data were continuously saved and backed up. All components were integrated with 10 Gigabit Ethernet (10G network) to provide a cohesive solution (Supplementary Information F). The average node-to-node transfer rate was on the order of 250-450 MB/s, including limitations of hard disk speed.

Imaging data were collected from the Nanozoomer and then automatically transferred to a data acquisition system. This step ensured uninterrupted scanning regardless of the limited disk space on the Nanozoomer system relative to the amount of data being acquired. The data acquisition system is the central repository for image pre-processing including image cropping, conversion, compression (Supplementary Information G).

The quality control (QC) service was applied to all stages of experimentation and image data flow in order to correct and improve the pipeline process organically. The experimental pipeline process information was recorded in an internal Laboratory Information Management System (LIMS). It supported the workflow by recording the detailed status of each experimental stage for each brain. Similarly, a separate online QC portal dictated all the image pre-processing stages (Figure 4). Through the LIMS and QC portal, it was possible to flag damaged sections to avoid unnecessary post-processing and identified the need to repeat a specific processing stage.

**Figure 4.**
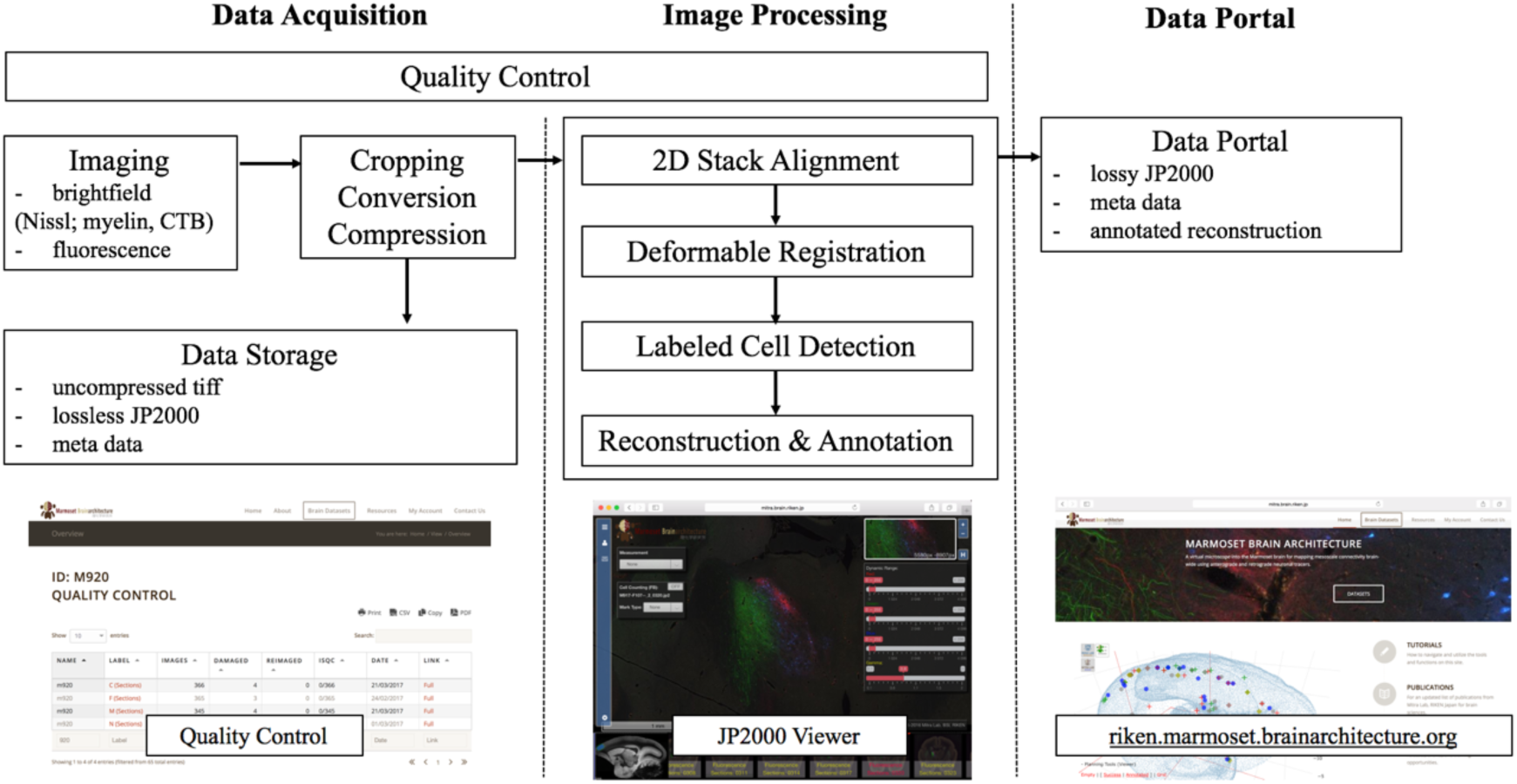
A flow chart showing the workflow of the computational pipeline, from data acquisition to image processing and finally dissemination on the public data portal. Arrows show the data flow. A Quality control system is implemented at every stage of the pipeline until final data release. The display of the data portals is to allow interactive service. (a) Quality control site (snapshots on the bottom left) which helps improve the pipelines process speed and manually flags unnecessary sections to avoid further post-processing. (b) An Openlayer 3.0 JPEG2000 viewer (snapshots on the bottom middle) including several controls such as dynamic range, gamma, measurement and auto cell detection tool to allow for a users’ interpretation. (c) The data portal site (snapshots on the bottom right) helps to host all successful and processed dataset for publishing purposes.

Image registration, cross-modal registration and automatic annotation, and tracing signal detection were performed in the image processing server. Images of individual sections were downsampled by 64 times and registered to one another using rigid-body transformation^52^. Registered 2D images were used to create a 3D volume of the brain in NIfTI format^53^ for each series. The transformation matrix for each downsampled image was applied to the corresponding full resolution image.

The brain outline of Brain/MINDs atlas^28^ was applied to the downsampled images after 2D registration to separate the brain regions from background and ventricles. Automatic annotation of the brain structures was achieved by registering the Brain/MINDs atlas to *ex-vivo* MRI and then aligned to the 2D registered Nissl series (“target images”). A 3D global affine transformation was applied to move the atlas images into the coordinate space of the MRI images. After transformation, the atlas images was matched to the MRI images using Large Deformation Diffeomorphic Metric Mapping (LDDMM)^54^ which transforms the atlas coordinate to the MRI image coordinate system. The same method was applied again to the transformed atlas images in order to match the target Nissl images. Individual brain regions could be automatically identified based on the transformed atlas. Figure 5a shows the example of automatic registration from Brain/MINDs atlas to target Nissl images. Cross-series registration using Euler2DTransform from Insight Segmentation and Registration Toolkit^55^ was performed to align 64-time downsampled myelin, CTB and fluorescence series of images to target Nissl images (Figure 5b-d). Finally, the transformation matrices calculated from the downsampled images were applied to the corresponding full resolution images. The annotations from the transformed atlas were aligned with the histology images of each series.

**Figure 5.**
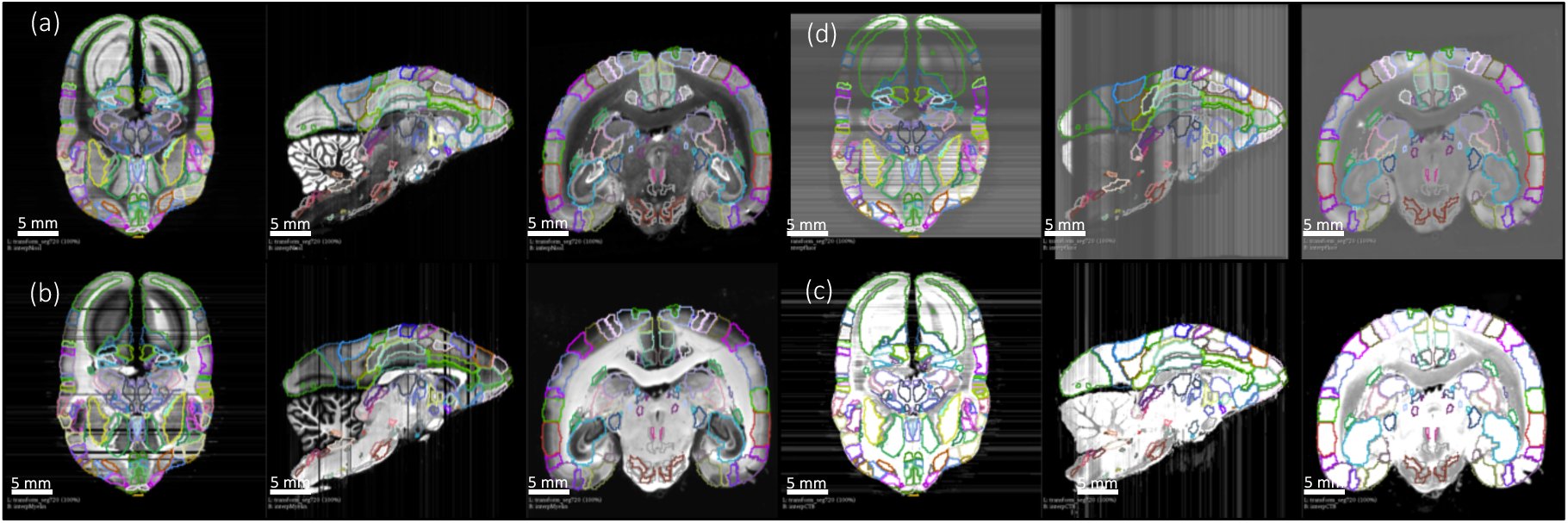
3D deformable registration and atlas mapping of all four series. The Brain/MINDs atlas was registered with *ex-vivo* MRI volume, and subsequently registered to target Nissl series (a). The shaded areas indicate missing sections at the end of processing (quality control). Other series including (b) myelin, (c) CTB and (d) fluorescence series were cross-registered to target Nissl series, and aligned with the atlas annotations. Only gray scale images are shown and they are sufficient for the registration process. Sample sections in transverse (left), sagittal (middle), and coronal (right) were shown for each series.

Injection volume was estimated by measuring the tracer spread at the injection site. Automatic cell and process detection was applied to individual registered sections in order to compute a draft whole-brain connectivity matrix. As an integral part of the computational pipeline, a data portal was developed to allow for viewing and interpreting high-resolution images online (http://riken.marmoset.brainarchitecture.org). By incorporating an Openlayer 3.0 image server with a custom image viewer, the data portal allows fully interactive zoom and pan, supports online adjustment of RGB dynamic range and contrast, as well as gamma adjustment (Figure 4). The data portal also provides visualization of cell detection results and an interactive tool for injection volume measurement.

#### Successful re-assembly of 3D volumes

In order to evaluate the quality of the image registration pipeline, we applied computational approaches to separately register series acquired for individual data modalities into separate volumes. Both high-quality and low-quality section images with staining issues, image variation, or artefacts were considered in the process. Adoption of the tape transfer method allowed us to maintain the geometry of the brain sections in the high-quality 20µm section images. This allowed successful section-to-section (2d) alignment using only rigid-body transformations. Poor-quality sections were selected by visual inspection and excluded from the 2d alignment step. Figure 6 (left) shows one marmoset brain with different staining procedures in coronal, sagittal and transverse planes after image reconstruction. It also shows the results of how the geometry of the brain has been maintained in each series.

**Figure 6.**
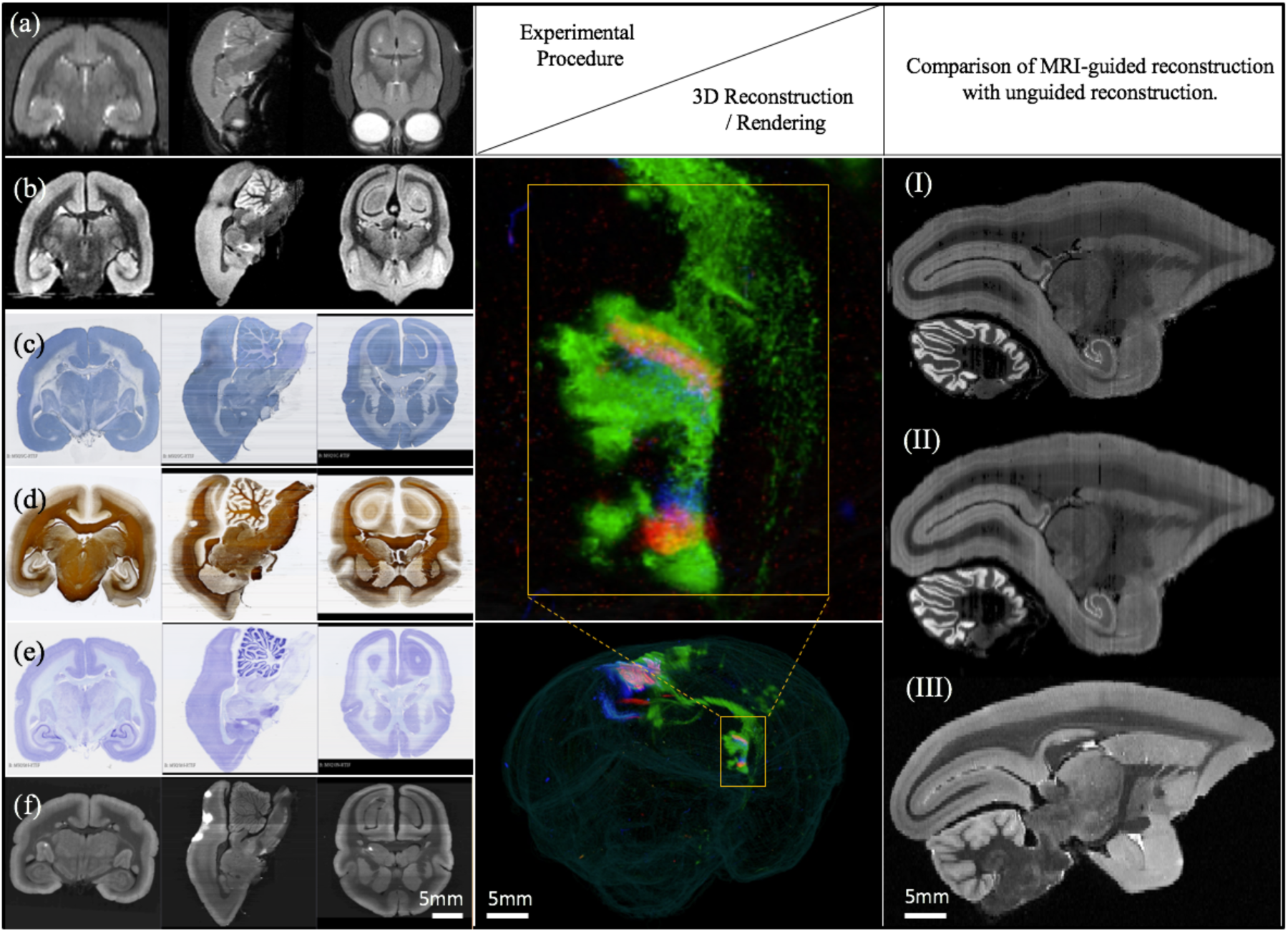
(left) Views of one marmoset brain after each experimental protocol: (a) *in-vivo* MRI (b) *ex-vivo* MRI (c) CTB staining (d) myelin staining (e) Nissl staining (f) fluorescence imaging. Coronal, sagittal and transverse planes at the same (MRI) or consecutive sections (staining series) are shown with 3D registration and reconstruction. (middle) A 3D visualization of the fluorescent tracer projection. Simultaneous anterograde (red, green) and retrograde (blue) tracing reveals a reciprocal connection between the dorsal medial visual area (injection site) and the thalamus (anterograde projection and retrograde cell labeled sites) especially lateral posterior nucleus and lateral pulvinar. The connectivity can be observed with this 3D visualization which shows the pathway of tracers in through the brain volume. (right) Comparison of MRI-guided reconstruction with unguided reconstruction. I: the target Nissl stack reconstruction by unguided piecewise neighbor-to-neighbor alignment. II: the MRI-guided reconstruction. III: same-subject T2-weighted MRI.

#### Atlas registration

Using external references such as the same-subject *ex-vivo* MRI or the population-typical reference atlas ^28^, we aimed to reconstruct the true shape of the subject brain and to avoid the classical curvature recoverability problem of sectioned objects. This atlas-informed reconstruction ^56^ improved reconstruction accuracy compared to the atlas-uninformed neighbor-to-neighbor method, as well as reduced the deformable metric cost. The impact of the *ex-vivo* MRI constraint on the 3D reconstruction is shown in Figure 6 (right). A visible distortion is present in the MRI-unguided reconstruction. This distortion is corrected by a MRI-guided method using a reference atlas. The MRI-constrained alignment of the Nissl sections produces a Nissl volume which closely resembles the convex hull of the same-subject MRI, leading to accurate parcellation of the brains in question.

## 3 Results

Brain volumes generated by the combined pipeline were further subjected to automated cross-modal registration and atlas segmentation, to obtain a regional connectivity matrix.

### Connectivity mapping

The registration process permitted brain surface reconstruction (Supplementary Video), 3D visualizations of projections, and virtual cuts in other planes of section than the original Coronal sections (Figure 6 (right)). After segmentation and registration, we derived quantitative values of tracer signals within each region. We developed an image processing method for detecting axonal and dendritic fragments in images, and applied it to each high resolution section (0.46µm) to segment the anterograde projections. The segmented pixels were appropriately weighted to create an isotropic 3D summary of the projections ^36^. We developed an automatic cell detection method ^57^ to segment somata labeled by the retrograde label Fast Blue throughout the entire brain. Injection sites were separated out from the rest of the brain. The projection strength between each target and source region was quantified as the fractional number of voxels containing tracer label.

The registration process together with process and cell detection methods allowed us to obtain intermediate resolution, annotated images for each tracer and to review the atlas parcellation. Figure 7 shows the result of three fluorescent tracer injections in the same animal and their origin/projections, resulting in one column and two rows in the putative connectivity matrix. In this example, Fast Blue, AAV-GFP and AAV-TdTOM were injected in V6, V1, and V6 visual cortex respectively. Automatic process detection identified projection targets from V1 to various regions, including the most prominent projections detected in V5 and dorsal lateral geniculate nucleus (DLG). Projection targets from V6 included the lateral pulvinar (LPul) and medial pulvinar (MPul) among other targets. Automatic cell detection for the Fast Blue tracer identified the regions projecting to V6 including prominent projections from A6DC, A31, and inferior pulvinar (IPul).

**Figure 7.**
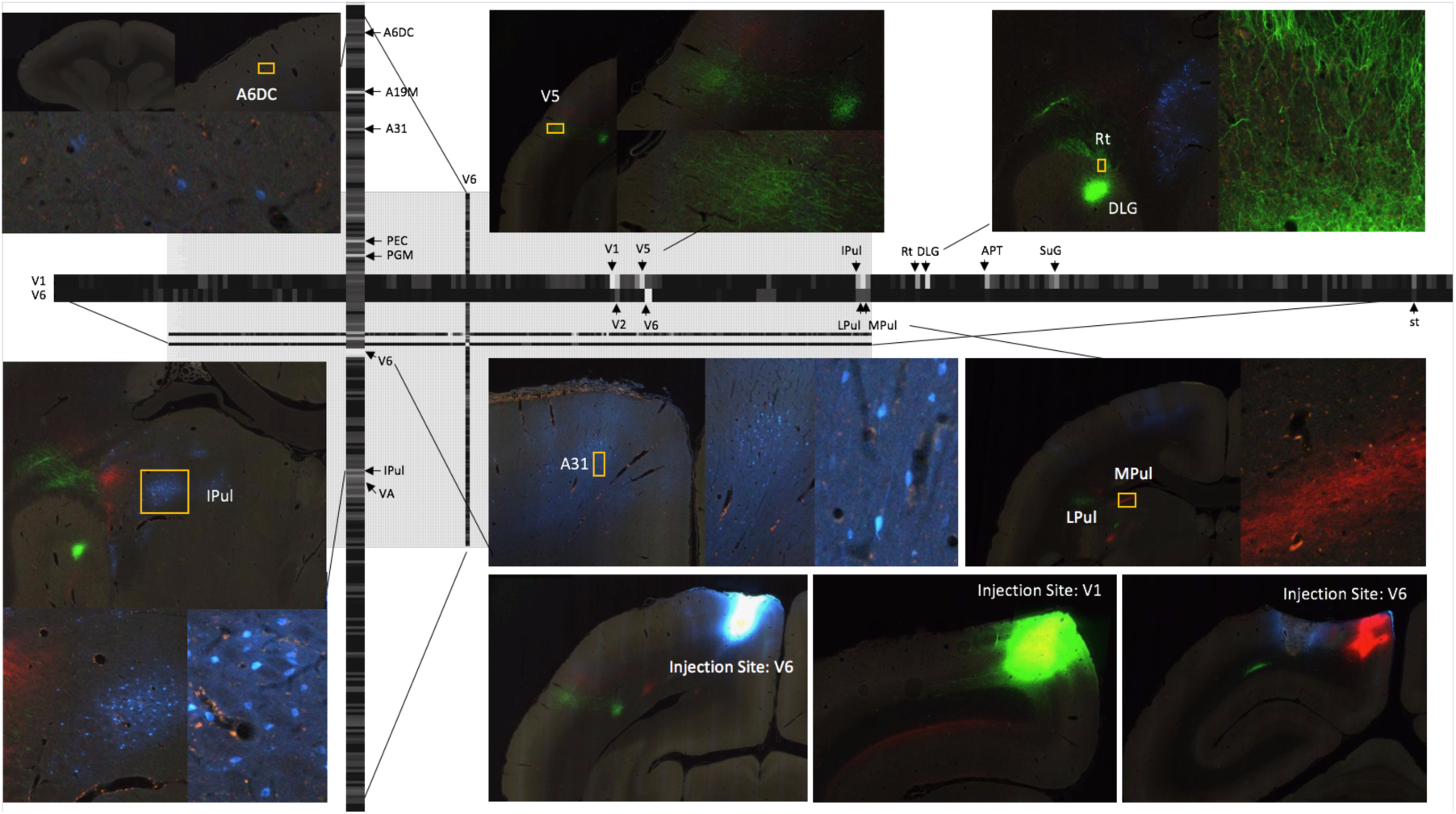
A part of the connectivity matrix identified with tracer injections in one sample brain. The retrograde tracer Fast Blue was injected in V6 and found in high density in several regions such as lPul and A31. AAV-GFP was injected in V1 and AAV-TdTOM in V6 and show clear projections to the thalamus and other visual areas. Each row contains all projections to different brain regions originating from those AAV tracers. The magnified images highlight some clear origin/projections from the injected tracers in the connectivity matrix.

## 4 Discussion

We have described a high throughput, standardized pipeline integrating experimental and computational elements into a unified system and workflow for processing tracer-injected Marmoset brains, representing an essential step towards producing a whole-brain mesoscale connectivity map in an NHP. The pipeline combines the well-established neuroanatomical protocols with automated instrumentation and a software system for greatly improving the efficiency of the techniques compared to conventional manually-intensive processing. Access to high-quality *in-vivo* and the *ex-vivo* MRI provided us with important auxiliary data sets facilitating re-assembly of the section images and atlas mapping, thus ameliorating the challenges arising from increased individual variations in brain geometry in an NHP compared with laboratory mice.

It is important to compare with other microscopic methods that have become established in recent years for light-microscope based anatomy, including serial block-face two photon scanning microscopy and light sheet microscopy, as well as knife-edge scanning microscopy. While these methods have important advantages, particularly the reduced need for section-to-section registration to produce the initial 3D volumes for further analysis, the classical methods have the important advantage of carrying through conventional histochemistry without major protocol alterations, producing long-lasting stains and precipitates that can be imaged using brightfield microscopy. Classical Nissl and myelin stains remain the gold standard for cytoarchitectonic texture-based determination of precise brain region location and delineation. These series are produced routinely with ease in the pipeline. The thin physical sections can be imaged rapidly in whole-slide imaging scanners rapidly and at relatively high numerical aperture (resolution in light sheet microscopy is comparatively limited due to reduced NA in the bulk of the sample).

The pipeline described here is for 1 ×; 3 glass slides that fortunately are large enough to accommodate Marmoset monkey brains in coronal section. The pipeline can be generalized in the future to 2 ×; 3 slides, which can handle larger brains (such as that of Macaque monkeys), with a few technical innovations, importantly stainers/coverslippers for the larger format slides. This should allow the easy and economical neurohistological processing of larger sized vertebrate brains, opening up the possibilities of applying modern computational neuroanatomical techniques to a significantly broader taxonomic range of species, allowing for the study of comparative neuroanatomical questions with unprecedented computational depth.

## 5 Acknowledgements

The authors would like to acknowledge Tetsuo Yamamori, Akiya Watakabe and Hiroaki Mizukami at RIKEN Center for Brain Science (CBS) for providing the virus tracers. We thank Noritaka Ichinohe and Yoko Yamaguchi at RIKEN CBS as well as Daniel Ferrante from CSHL Mitra Lab for helping with computational aspects of the pipeline. We acknowledge the effort from our previous technicians, Kevin Weber and Khurshida Hossain (RIKEN) and the support from Erika Sasaki at the Central Institute for

Experimental Animals for providing marmosets used in this project. We thank staff of the Biomedicine Discovery Institute, (Monash University) for providing a detailed protocol for injection surgeries (Katrina Worthy), helping perform surgeries (Jonathan Chan), and providing helpful insight on the development of the web infrastructure (Shi Bai and Piotr Majka). We acknowledge James Bourne, Inaki-Carril Mundinano and William Kwan (Australian Regenerative Medicine Institute, Monash University), who performed MRI-guided stereotaxic brain injections in deep brain structures. We acknowledge the support from Jaikishan Jayakumar at IIT Madras on neuroanatomical questions, as well as Xu Li from CSHL machine vision algorithms for process detection. This project is supported by the Brain Mapping of Integrated Neurotechnologies for Disease Studies (Brain/MINDS) from the Japan Agency for Medical Research and Development, AMED under grant number JP17dm0207001, the Crick-Clay Professorship (CSHL), the Mathers Foundation and the H N Mahabala Chair at IIT Madras.

